# A Superfamily of T6SS Antibacterial Effectors Displaying L,D-carboxypeptidase Activity Towards Peptidoglycan

**DOI:** 10.1101/2020.02.18.954545

**Authors:** Stephanie Sibinelli de Sousa, Julia Takuno Hespanhol, Gianlucca Gonçalves Nicastro, Bruno Yasui Matsuyama, Stephane Mesnage, Ankur Patel, Robson Francisco de Souza, Cristiane Rodrigues Guzzo, Ethel Bayer-Santos

## Abstract

Type VI secretion systems (T6SSs) are contractile nanomachines used by bacteria to inject toxic effectors into competitors. The identity and mechanism of many effectors remain unknown. We characterized a *Salmonella* SPI-6 T6SS antibacterial effector called Tae5^STM^ (<u>t</u>ype VI <u>a</u>midase <u>e</u>ffector 5). Tae5^STM^ is toxic in target-cell periplasm and is neutralized by its cognate immunity protein (Tai5^STM^). Microscopy analysis revealed that cells expressing the effector stop dividing and lose cell envelope integrity. Bioinformatic analysis uncovered similarities between Tae5^STM^ and the catalytic domain of L,D-transpeptidase. Point mutations on conserved catalytic histidine and cysteine residues abrogated toxicity. Biochemical assays revealed that Tae5^STM^ displays L,D-carboxypeptidase activity, cleaving peptidoglycan tetrapeptides between *meso*-diaminopimelic acid^3^ and D-alanine^4^. Phylogenetic analysis showed that Tae5^STM^ homologs constitutes a new superfamily of T6SS-associated amidase effectors distributed among α-, β- and γ-proteobacteria. This work expands our current knowledge about bacterial effectors used in interbacterial competition.

## Introduction

Bacteria commonly live in densely populated polymicrobial communities and compete over scarce resources. Several types of contact-dependent antagonistic interactions between bacteria have been described (Garcia-Bayona and Comstock, 2018). The type VI secretion system (T6SS) is a dynamic contractile structure evolutionarily related to bacteriophage tails that delivers protein effectors in a contact-dependent manner into diverse cellular types, including eukaryotic host cells and rival bacteria and fungi (Hachani et al., 2016; Coulthurst, 2019; Trunk et al., 2019). T6SSs were also reported to display contactin-dependent functions in which secreted effectors facilitate the scavenging of scarce metal ions (Wang et al., 2015; Si et al., 2017a; Si et al., 2017b; DeShazer, 2019).

The T6SS is anchored in the bacterial envelope and is composed of 13 core structural components that assemble into three major complexes: the trans-membrane complex, the baseplate and the tail (Nguyen et al., 2018). A conformational change in the T6SS baseplate is thought to trigger the contraction of a cytoplasmic sheath, expelling a spear-like structure to puncture target cell membrane (Wang et al., 2017; Salih et al., 2018). The spear is composed of Hcp (hemolysin co-regulated protein) hexamers capped with a trimer of VgrG (valine-glycine repeat protein G) proteins and a PAAR (proline-alanine-alanine-arginine) protein tip (Mougous et al., 2006; Renault et al., 2018; Shneider et al., 2013). Cargo effector proteins associate through non-covalent interactions with these structural components, while specialized effectors are presented as additional C-terminal domains fused to Hcp, VgrG, or PAAR protein (Cianfanelli et al., 2016; Jana and Salomon, 2019). Consequently, along with the Hcp-VgrG-PAAR puncturing device, a cocktail of effectors is delivered into the target cell after each contraction event.

Antibacterial effectors delivered by the T6SS induce toxicity by targeting important structural components or affecting target-cell metabolism. Several families of effectors have been described, including peptidoglycan amidases and hydrolases, phospholipases, nucleases, NAD(P)^+^-glycohydrolases, pore forming proteins and enzymes that synthesize (p)ppApp (Russell et al., 2012; Koskiniemi et al., 2013; Whitney et al., 2013; Ma et al., 2014; Tang et al., 2018; Ahmad et al., 2019; Jana et al., 2019; Mariano et al., 2019; Wood et al., 2019). An example of an effector inducing toxicity by posttranslational modification (ADP-ribosylation) of the cytoskeleton component FtsZ has also been reported (Ting et al., 2018). In order to prevent self-intoxication, bacteria have a specific immunity protein for each antibacterial T6SS effector. Immunity proteins are encoded adjacent to their cognate effector, reside in the same cellular compartment where the effector exerts its toxic effect, and typically work by binding directly to the effector (Hood et al., 2010; Russell et al., 2012).

The peptidoglycan sacculus maintains cell shape and provides mechanical strength to resist osmotic pressure (Vollmer et al., 2008). The mesh-like structure surrounds the cytoplasm and inner membrane and is composed of glycan chains of alternating *N*-acetylglucosamine (NAG) and *N*-acetylmuramic acid (NAM) residues crosslinked by short peptides containing both L- and D-amino acids. In Gram-negative bacteria, the peptide component is usually made of the following amino acids: L-alanine^1^, D-*iso*glutamic acid^2^ (D-*i*Glu), *meso*-diaminopimelic acid^3^ (*m*DAP), D-alanine^4^ and D-alanine^5^ (Typas et al., 2011). Peptide stems are crosslinked to each other by transpeptidases that could be either D,D-transpeptidases, forming crosslinks between D-Ala^4^ of a pentapeptide donor stem and *m*DAP^3^ of a tetrapeptide acceptor stem (4 → 3 crosslink); or L,D-transpeptidases, forming crosslinks between *m*DAP of one tetrapeptide stem and *m*DAP of another tetrapeptide stem (3 → 3 crosslink) (Vollmer and Bertsche, 2008). D,D-transpeptidases (also called penicillin-binding proteins, PBPs) are the primary enzymes performing crosslinks in the peptidoglycan and are sensible to beta-lactam antibiotics, which mimic the terminal D-Ala^4^-D-Ala^5^ moiety of the donor pentapeptide (King et al., 2016).

T6SS effectors targeting the peptidoglycan distribute into two groups: 1) those that act as amidases and cleave within the peptide stems or crosslinks (Russell et al., 2012); and 2) those that act as glycoside hydrolases and cleave the glycan backbone (Whitney et al., 2013). T6SS amidase effectors form four phylogenetically distinct families named Tae1-4 (<u>t</u>ype VI <u>a</u>midase <u>e</u>ffectors) and the preferred cleavage site within the peptidoglycan varies between each family (Russell et al., 2012). Tae1 and Tae4 cleave the bond between D-*i*Glu^2^ and *m*DAP^3^ within the same peptide stem, while Tae2 and Tae3 cleave the crosslink bridge between D-Ala^4^ and *m*DAP^3^ of different peptide stems (Russell et al., 2012). An effector named TaeX was recently added to the list of Tae effectors and its only representative cleaves the amide bond between NAM glycan and the first L-Ala^1^ of the peptide stem (Ma et al., 2018). The superfamily of T6SS glycoside hydrolases was divided into three families named Tge1-3 (<u>t</u>ype VI <u>g</u>lycoside hydrolase <u>e</u>ffector) (Whitney et al., 2013). Tge members have a lysozyme-like fold and were shown to cleave the glycoside bond between NAM and NAG (Whitney et al., 2013).

In *Salmonella enterica*, T6SS gene clusters are encoded within different pathogenicity islands (SPIs), depending on the subspecies and serovar (Blondel et al., 2009; Bao et al., 2019). *S. enterica* subsp. *enterica* serovar Typhimurium (*S*. Typhimurium) encodes a T6SS within SPI-6. The expression of SPI-6 T6SS genes is not detected under laboratory culture conditions, but it is activated in later stages of macrophage infection (Parsons and Heffron, 2005; Mulder et al., 2012). In addition, SPI-6 T6SS is active in the mammalian gut where it works as an antibacterial weapon to kill the resident species of the microbiota and contribute to *Salmonella* colonization (Sana et al., 2016). The histone-like nucleoid structuring protein (H-NS) (Brunet et al., 2015) and the ferric uptake regulator (Fur) (Wang et al., 2019) were reported to repress SPI-6 T6SS genes *in vitro*. Although the SPI-6 T6SS was described to be involved in bacterial replication inside macrophages (Mulder et al., 2012) and in interbacterial competition inside the gut (Sana et al., 2016), only one effector (Tae4) was described to date as a substrate for SPI-6 T6SS. Tae4 is an antibacterial effector and works as a D,L-endopeptidase cleaving the bond between D-*i*Glu^2^ and *m*DAP^3^ (Russell et al., 2012; Benz et al., 2013; Zhang et al., 2013).

Here, we set out to identify new SPI-6 T6SS effectors and report the characterization of a new superfamily of T6SS antibacterial effectors containing the domain of unknown function DUF2778, which displays L,D-carboxypeptidase activity and cleaves peptidoglycan tetrapeptides between *m*DAP^3^ and D-Ala^4^ within the same peptide stem. This superfamily is evolutionarily related to enzymes that have an L,D-transpeptidase fold and is broadly distributed among α-, β- and γ-proteobacteria. We also describe a protein containing DUF2195 as a cognate immunity protein for Tae5^STM^. Expression of Tae5^STM^ in the periplasm of target *Escherichia coli* cells prevents cell division, induces cell elongation, swelling and lysis. Our data suggests that the L,D-carboxypeptidase activity of Tae5^STM^ preferentially targets the acceptor tetrapeptide stem, thus preventing formation of new crosslinks by D,D-transpeptidases during peptidoglycan synthesis and resulting in altered formation of the division septum and an overall weakened peptidoglycan structure.

## Results

### Tae5^STM^/Tai5^STM^ are a new antibacterial effector/immunity pair

To search for new T6SS effectors secreted by *S*. Typhimurium we inspected the SPI-6 T6SS gene cluster of the 14028s strain looking for bicistrons that could resemble an antibacterial effector/immunity pair (Figure 1A). The gene annotated as *STM14_0336* encodes a small protein of 173 amino acids and contains a DUF2778 (PF10908). Upstream to this gene, there is another small protein (138 amino acids) containing a DUF2195 annotated as *STM14_0335*, which encodes a predicted Sec signal peptide sequence for periplasmic localization (Figure S1A) (Almagro Armenteros et al., 2019). Bastion6 software prediction (Wang et al., 2018) indicates that STM14_0336 could be a T6SS effector (score 0.758). To test whether *STM14_0336* and *STM14_0335* are a bona fide effector/immunity pair we cloned these genes in compatible vectors under the control of different promoters. To evaluate the toxicity of STM14_0336 upon expression in *E. coli* and to establish in which cellular compartment the toxin exerts its effect, *STM14_0336* was cloned into pBRA vector under the control of Pbad promoter (inducible by L-arabinose and repressed by D-glucose) both with and without an N-terminal PelB periplasmic localization sequence (pBRA SP-Tae5^STM^ or pBRA Tae5^STM^). The putative STM14_0335 immunity protein was cloned with its endogenous signal peptide into pEXT22 vector under the control of P_TAC_ promoter, which can be induced by IPTG (pEXT22 Tai5^STM^). *E. coli* strains carrying different combinations of pBRA and pEXT22 plasmids were serially diluted and incubated on LB-agar containing either 0.2% D-glucose or 0.2% L-arabinose plus 200 μM IPTG (Figure 1B). Results showed that STM14_0336 is toxic when directed to the periplasm of *E. coli* (pBRA SP-Tae5^STM^) but not in the cytoplasm (pBRA Tae5^STM^), and that STM14_0335 (pEXT22 Tai5^STM^) could neutralize STM14_0336 toxicity (Figure 1B). To confirm the subcellular localization of the immunity protein, *E. coli* cells expressing a C-terminal FLAG-tagged version of STM14_0335 were subjected to subcellular fractionation and results confirmed its periplasmic localization (Figure 1C).

**Figure 1.**
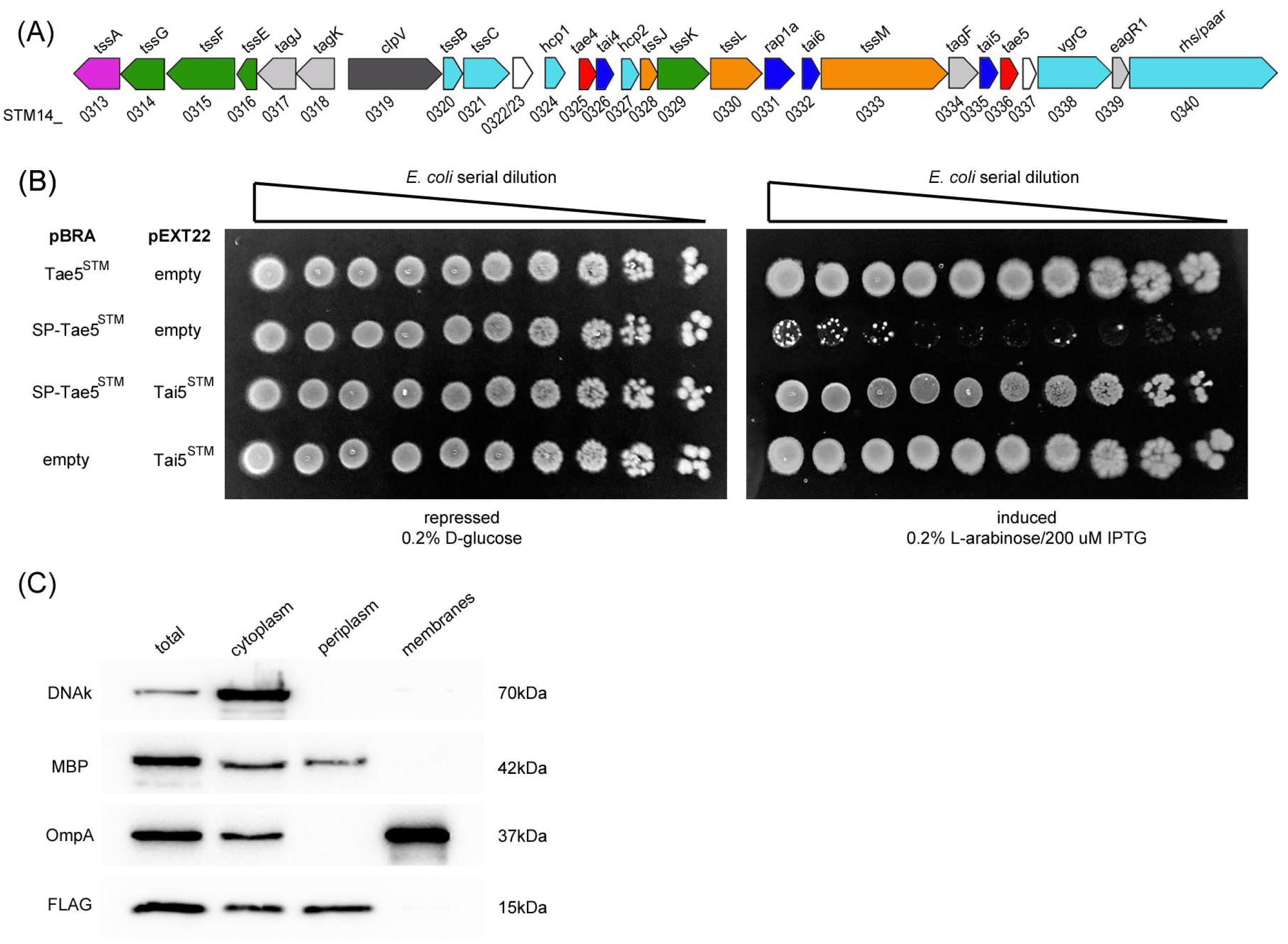
Tae5^STM^/Tai5^STM^ are a new antibacterial effector/immunity pair. (A) Schematic representation of SPI-6 T6SS gene cluster of *S*. Typhimurium 14028s: membrane complex (orange), baseplate (green), tail (light blue), toxins (red), immunity proteins (dark blue), chaperone and stabilizing protein (pink), ATPase for disassembly (dark gray), accessory (light gray). KEGG accession numbers are indicated below. (B) Four-fold dilutions of *E. coli* strains containing pBRA and pEXT22 constructs as indicated spotted onto LB-agar plates. Growth inhibition is observed upon expression of the SP-Tae5^STM^ construct and can be neutralized by co-expression of Tai5^STM^. (C) *E. coli* cells expressing pEXT22 Tai5^STM^-FLAG were fractionated and analyzed by western blot using anti-FLAG, anti-DNAk (cytoplasmic), anti-OmpA (membranes) and anti-MBP (periplasm) antibodies.

### Tae5^STM^ intoxication causes altered cell division, swelling and lysis

To gather further insight on the mechanism by which Tae5^STM^ induce toxicity, we performed time-lapse microscopy to evaluate growth and morphology of individual *E. coli* cells carrying pBRA SP-Tae5^STM^. These *E. coli* cells grew normally when incubated on LB-agar pads containing 0.2% D-glucose (repressed) over a timeframe of 16h (Figure 2A, Movie S1). However, incubation in the presence of 0.2% L-arabinose induced a series of alterations in cell division and morphology (Figures 2A-F, Movie S2). At early time points (up to 4.5h) after induction with L-arabinose, intoxicated cells tend to stop dividing or divide with increased doubling times (Figures 2B and 2C). Between 0 and 4.5h of incubation in L-arabinose, only 52% of intoxicated cells were observed to undergo at least one round of cell division with a doubling time of 93 ± 34 min, while 84% of cells grown with D-glucose divided in the same timeframe with a doubling time of 54 ± 20 min (Figures 2B and 2C). Although intoxicated cells had clearly impaired cell division, they continued to increase in size (Figure 2D, Movie S2). After 4.5h, cells grown in L-arabinose had an average cell length of 20 ± 12 μm, while cells grown in D-glucose displayed an average length of 3.7 ± 0.7 μm (Figure 2D). To analyze whether intoxicated cells were not dividing or dividing but not segregating into daughter cells, we incubated *E. coli* with the membrane dye FM 4-64 (Figure 2E). Results showed that elongated cells lack membrane invaginations indicative of a division septum (Figure 2E). To access if the assembly of the division septum was perturbed, we transformed an *E. coli* strain in which mVenus was fused to the endogenous *ftsZ* gene (Moore et al., 2017) with our pBRA SP-Tae5^STM^ plasmid (Figure 2F). Data show that the Z-ring formed by FtsZ-mVenus was not properly assembled in intoxicated cells, with FtsZ labelling dispersed throughout the cell (Figure 2F).

**Figure 2.**
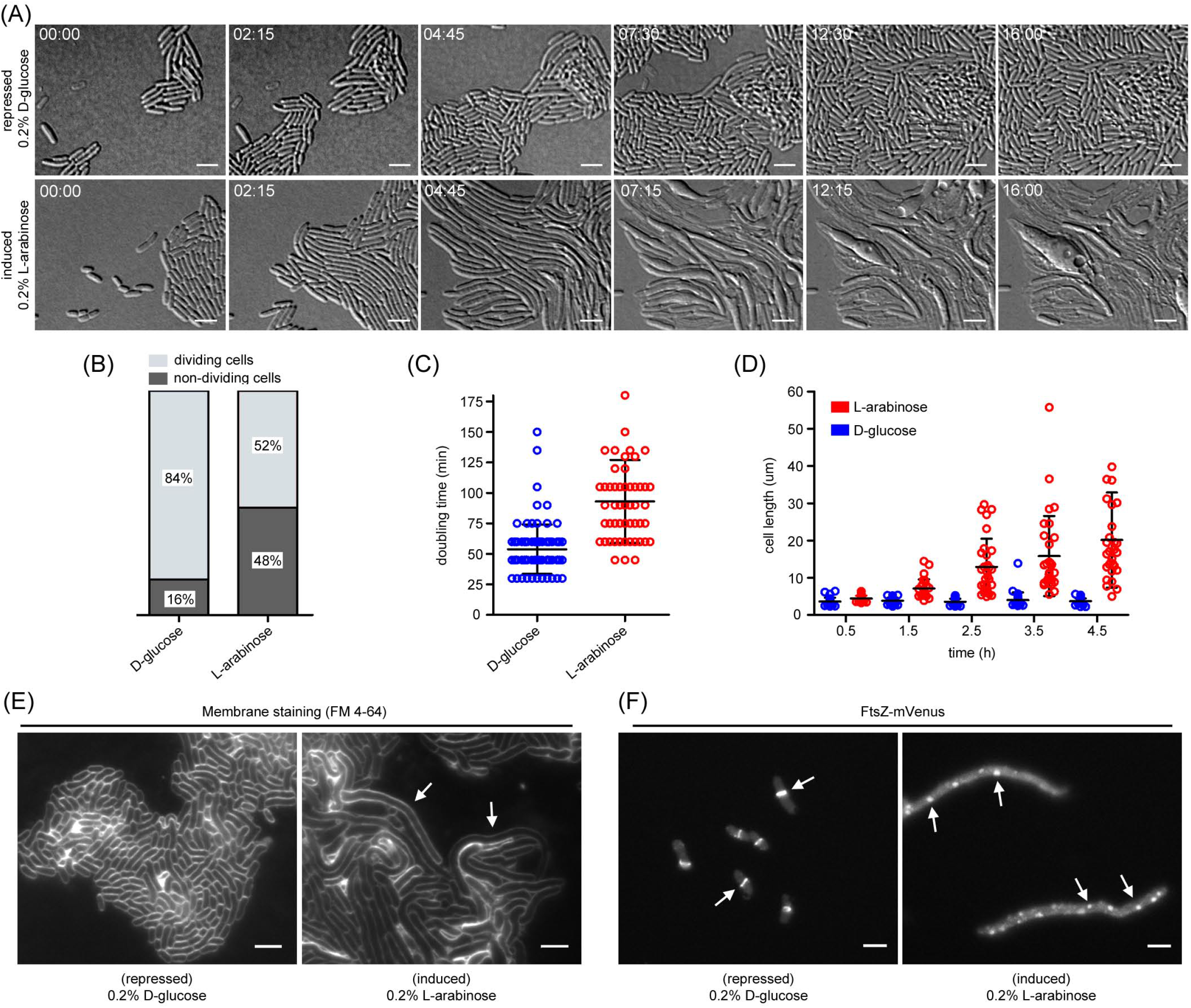
Tae5^STM^ is a periplasmic-acting toxin that alters cell division and weakens the peptidoglycan. (A) Time-lapse microscopy of *E. coli* cells expressing SP-Tae5^STM^ grown on LB-agar pads containing either 0.2% D-glucose (repressed) or 0.2% L-arabinose (induced). Scale bar 5 μm. Timestamps in hours:minutes. (B) Percentage of dividing and non-dividing cells observed in (A) between 0 and 4.5h. (C) Doubling time in minutes of cells that divided quantified in (B). (D) Cell length of cells observed in (A) between 0 and 4.5h. (E) Fluorescence microscopy images of *E. coli* cells harboring pBRA SP-Tae5^STM^ labeled with the membrane dye FM 4-64 and incubated with 0.2% D-glucose (repressed) or 0.2% L-arabinose (induced). Arrows indicate elongated cells with no septum membrane invagination. Scale bar 5 μm. (F) Fluorescence microscopy images of *E. coli* FtsZ-mVenus carrying pBRA SP-Tae5^STM^ incubated with 0.2% D-glucose (repressed) or 0.2% L-arabinose (induced). Arrows indicate FtsZ labelling corresponding to the Z-ring (left panel) or dispersed throughout the cell (right panel). Scale bar 5 μm.

After 4.5h in L-arabinose, *E. coli* pBRA SP-Tae5^STM^ cells tend to swell and finally burst, indicating that the integrity of their peptidoglycan structure was compromised (Figure 2A, Movie S2). No obvious alterations were observed in *E. coli* pBRA SP-Tae5^STM^ cells grown in the presence of D-glucose during the same timeframe (Figure 2A, Movie S1).

### Tae5^STM^ is evolutionarily related to L,D-transpeptidases but display L,D-carboxypeptidase activity

To gain insight into the molecular function of Tae5^STM^, we used its amino acid sequence (STM14_0336) as query in JackHMMER searches (Potter et al., 2018) to fetch a total of 143242 sequences with significant similarity (inclusion threshold ≤ 10^-6^ and e-value ≤ 10^-3^) from the NCBI nr database (June 7^th^, 2019). Additional JackHMMER searches using selected hits of the first iterative search as queries and the Pfam models DUF2778 (PF10908), YkuD (PF03734) and YkuD_2 (PF13645), resulted in a total of 153327 sequences. To reduce data complexity, we clustered amino acid sequences requiring 80% coverage for all pairwise alignments generated by MMseqs (Steinegger and Soding, 2017), and a maximum e-value of 10^-3^, resulting in 4113 groups. A single amino acid sequence was chosen as representative from each group with at least 9 members, resulting in 943 sequences representing a sample of 145969 homologs. These representative sequences were used to build a phylogenetic tree using the maximum likelihood method (Figure 3A). Homologs of Tae5^STM^ clustered into 5 main clades: clade 1 (red, 6332 sequences), to which Tae5^STM^ belongs, is composed of proteins containing the uncharacterized DUF2778 domain (DUF2778 superfamily); clade 2 (blue, 44540 sequences) contains the L,D-transpeptidases from *Bacillus subtilis* (Ldt_Bs_, PDB 1Y7M) (Bielnicki et al., 2006); clade 3 (green, 44090 sequences) contains L,D-transpeptidases from *Enterococcus faecium* (Ldt_fm_, PDB 1ZAT) (Biarrotte-Sorin et al., 2006), *Mycobacterium abscessus* (LdtMa_b_, PDB 5UWV) (Kumar et al., 2017) and *Mycobacterium tuberculosis* (Ldt_Mt1-3_, PDB 3TUR, 5DCC) (Erdemli et al., 2012; Bianchet et al., 2017); clade 4 (purple, 25242 sequences) contains an enzyme from *Helicobacter pylori* (Cds6, PDB 4XZZ) that has a catalytic domain resembling L,D-transpeptidases but with L,D-carboxypeptidase activity (Kim et al., 2015); clade 5 (gray, 25765 sequences) contains an L,D-transpeptidase from *E. coli* (YcbB, PDB 6NTW) (Caveney et al., 2019) and proteins recognized by the Pfam model YkuD_2.

**Figure 3.**
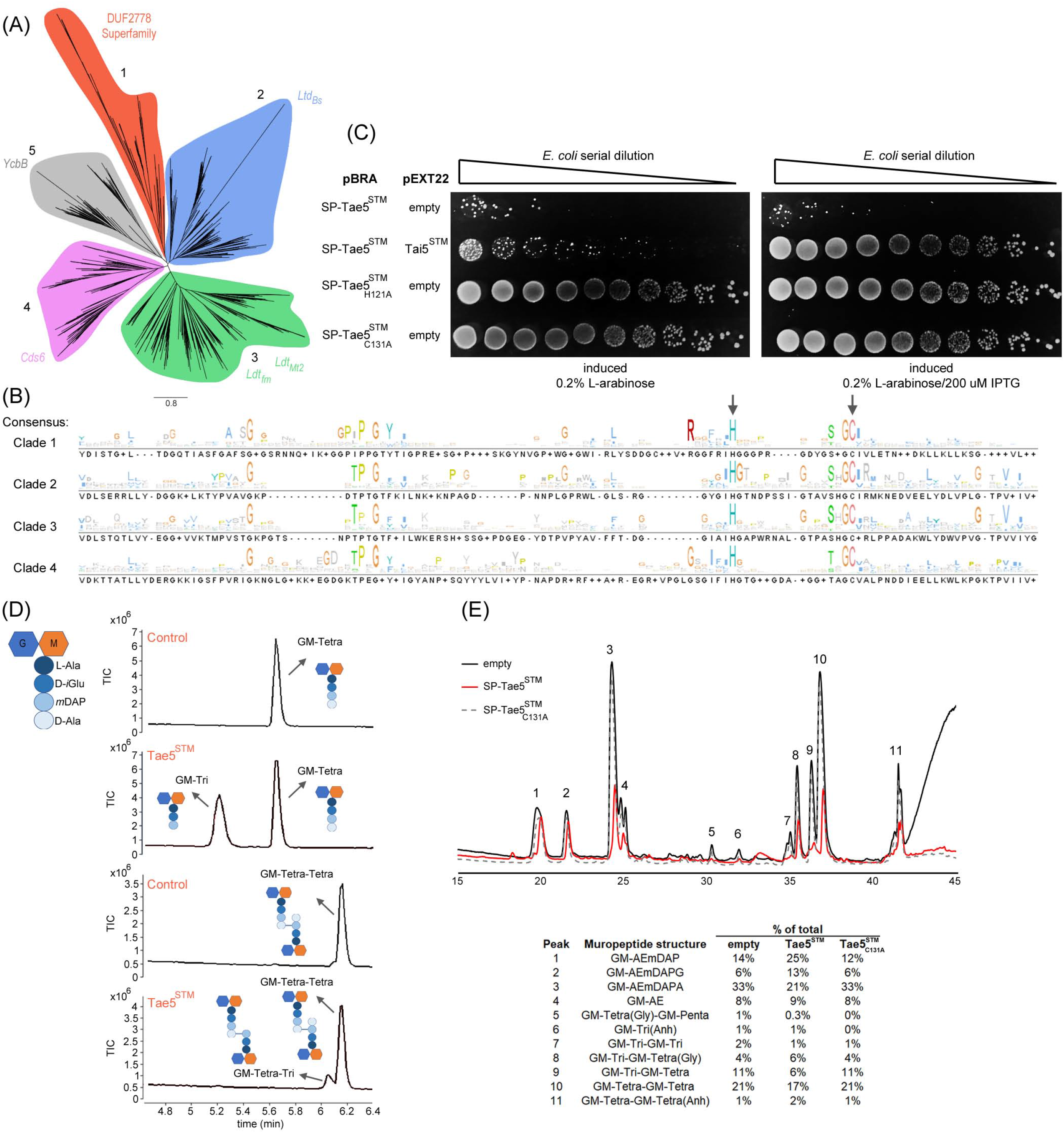
Tae5^STM^ is evolutionarily related to L,D-transpeptidases but display L,D-carboxypeptidase activity. (A) Maximum likelihood phylogenetic tree of DUF2778 homologs identified using JackHMMER and HMMsearch. Tae5^STM^ is a distant homolog of L,D-transpeptidases with YkuD domain (PF03734). DUF2778-containing proteins grouped separately (clade 1, red) from known L,D-transpeptidases. (B) Partial amino acid sequence alignment of consensus sequence from clades 1-4. Arrows indicate conserved catalytic histidine and cysteine residues of L,D-transpeptidases. (C) Four-fold dilutions of *E. coli* strains containing pBRA and pEXT22 constructs as indicated spotted onto LB-agar plates. Growth inhibition is observed upon expression of SP-Tae5^STM^, but toxicity is abolished by H121A and C131A point mutations. (D) RP-HPLC coupled to MS showing purified monomeric GM-tetrapeptides or dimeric GM-tetrapeptide-GM-tetrapeptide incubated with recombinant Tae5^STM^. Schematic structure of muropeptides are shown. N-acetylglucosamine (G), N-acetylmuramic acid (M), L-alanine (L-Ala), D-*iso*glutamic acid (D-*i*Glu), *meso-* diaminopimelic acid (*m*DAP), D-alanine (D-Ala). (E) RP-HPLC coupled to MS showing total ion chromatograms of muropeptides obtained after mutanolysin digestion of peptidoglycan extracted from *E. coli* harboring empty pBRA (black), pBRA SP-Tae5^STM^ (red) and pBRA SP-Tae5^STM^_C131A_ (gray). Inferred muropeptides structures and relative abundance (% of total) of each peak quantified by MassHunter software (bottom).

Multiple amino acid sequence alignments of proteins from each clade revealed conserved residues similar to the conserved motif described for L,D-transpeptidases: <u>H</u>XX_14-17_[S/T]HG<u>C</u>h (underlined letters are absolutely conserved catalytic residues and “h” hydrophobic residues) (Erdemli et al., 2012) (Figure 3B). Interestingly, according to Pfam HMM logo and our sequence alignment (Figure 3B), the number of residues between the conserved catalytic His and Cys is smaller in DUF2778 (PF10908) compared to YkuD (PF3734). To evaluate whether the conserved His in position 121 and Cys in position 131 of STM14_0336 are required to induce the toxic phenotype, we produced point mutations by substituting these residues for alanine. Plasmids containing point mutations (pBRA SP-Tae5^STM^_H121A_ and pBRA SP-Tae5^STM^_C131A_) were transformed into *E. coli* cells and grown in the presence of 0.2% L-arabinose, revealing complete loss of toxicity (Figure 3C).

To determine the enzymatic activity of Tae5^STM^ we incubated purified recombinant protein (Figure S2A) with purified peptidoglycan tetrapeptides and analyzed the reaction product by reverse phase HPLC coupled to mass-spectrometry (Figure 3D). Results showed that Tae5^STM^ has L,D-carboxypeptidase activity and cleaves NAG-NAM-tetrapeptides (GM-tetrapeptide) between *m*DAP^3^ and D-Ala^4^, producing GM-tripeptides (Figure 3D). The formation of crosslinked GM-tripeptide-GM-tetrapeptide could not be detected, suggesting that Tae5^STM^ does not have L,D-transpeptidase activity (Figure 3D). Crosslinked dimeric GM-tetrapeptide-GM-tetrapeptide forms were also provided as substrate and a small proportion of cleavage was detected between the *m*DAP^3^-D-Ala^4^ of the acceptor peptide stem during the same incubation period, thus confirming the L,D-carboxypeptidase activity and suggesting that Tae5^STM^ preferentially uses monomeric GM-tetrapeptide as substrate (Figure 3D). Also, we could not detect any product indicating that Tae5^STM^ cleaves the D-Ala^4^-*m*DAP^3^ crosslink bridge between peptide stems (Figure 3D). Moreover, co-incubation of GM-tetrapeptides (GM-AE*m*DAP<u>A</u>) with an excess of D-methionine (1 mM) and recombinant Tae5^STM^ produced GM-tetrapeptides with D-Met at position 4 (GM-AE*m*DAP<u>M</u>), showing that Tae5^STM^ is able to exchange the last amino acid of GM-tetrapeptides (Figure S2B) as was reported for other L,D-transpeptidases (Mainardi et al., 2005).

The composition of peptidoglycan extracted from *E. coli* cells growing at exponential phase is enriched in monomeric GM-tetrapeptide and dimeric GM-tetrapeptide-GM-tetrapeptide (Glauner et al., 1988). During synthesis of new peptidoglycan, the GM-pentapeptide precursor works as donor peptide stem and is crosslinked to acceptor GM-tetrapeptide stems by the action of D,D-transpeptidases, forming 4 → 3 crosslink bridges (Typas et al., 2011). As Tae5^STM^ seems to preferentially degrade GM-tetrapeptides into GM-tripeptides *in vitro*, we hypothesized that the effector would promote toxicity by depleting the pool of acceptor GM-tetrapeptides, thus preventing the formation of new crosslinks by endogenous D,D-transpeptidases. To test this hypothesis, we extracted peptidoglycan from *E. coli* cells carrying 1) empty pBRA plasmid, 2) pBRA SP-Tae5^STM^ or 3) pBRA SP-Tae5^STM^_c131A_ incubated with 0.2% L-arabinose for 3h and analyzed the profile of muropeptides by HPLC-MS (Figure 3E). The muropeptide profile of *E. coli* cells expressing SP-Tae5^STM^ was reduced in GM-tetrapeptide (GM-AE*m*DAPA) (21%) compared to cells with empty plasmid (33%) or SP-Tae5^STM^_C131A_ (33%) (Figure 3E). Likewise, the proportion of GM-tripeptides (GM-AE*m*DAP) was enriched in cells expressing SP-Tae5^STM^ (25%) compared to empty and SP-Tae5^STM^_C131A_ (13%) (Figure 3E). Moreover, the proportion of most crosslinked forms were reduced in *E. coli* cells expressing SP-Tae5^STM^ compared to cells carrying the empty plasmid or expressing SP-Tae5^STM^_C131A_ (Figure 3E).

### DUF2778 superfamily is widespread and segregates into three families

Phylogenetic analysis of sequences from clade 1 comprising DUF2778 superfamily (Figure 3A) showed that proteins segregate into three families (Figure 4A). Given that Tae5^STM^ displays amidase activity (L,D-carboxypeptidase), we propose to rename its family Tae5 (<u>t</u>ype VI <u>a</u>midase <u>e</u>ffector 5) in line with current T6SS effector nomenclature (Russell et al., 2012; Whitney et al., 2013), and the other two families Tae6 and Tae7 (Figure 4A). Tae5 family contains the Tae5^STM^ effector and is composed mainly of *Salmonella enterica* subsp. *enterica* serovars encoding SPI-6 T6SS (γ-Proteobacteria, Enterobacteriaceae), followed by species of *Bordetella* (β-Proteobacteria, Burkholderiaceae) and some α-Proteobacteria from Rhizobiaceae and Sphingomonadaceae (Figure 4B and Table S1A). A few examples of *S. enterica* subsp. *diarizonae* (SPI-20 and SPI-21 T6SSs) and subsp*. salamae* (SPI-19 T6SS) were detected in Tae5 family (Figure 4B and Table S1A). In addition, Tae5 family also contains examples of Actinobacteria (Streptomycetaceae) and Cyanobacteria (Synechococcaceae, Microcystaceae, Aphanothecaceae); however, it is not clear at this point whether these proteins are secreted by alternative secretion mechanisms, such as the extracellular contractile injection systems (eCIS) (Chen et al., 2019), or simply work in the remodeling of peptidoglycan. Tae6 family is abundant in a large variety of species from Enterobacteriaceae *(Cedecea, Citrobacter, Cronobacter, Enterobacter, Escherichia, Franconibacter, Gibbsiella, Klebsiella, Kluyvera, Leclercia, Lelliottia, Salmonella)*, as well as Erwiniaceae *(Pantoea, Erwinia)* and Yersiniaceae *(Nissabacter, Rahnella, Yersinia)*. Examples of *S. enterica* subsp*. arizonae* (SPI-20 and SPI-21 T6SSs) and subsp*. diarizonae* (SPI-20 and SPI-21 T6SSs), and a few *S. enterica* subsp*. enterica* serovars such as Fresno (SPI-6 T6SS) were detected in Tae6 family (Figure 4B and Table S1B). Tae7 family is the most widespread and can be found in Enterobacteriaceae *(Cedecea, Citrobacter, Cronobacter, Enterobacter, Escherichia, Klebsiella, Kluyvera, Leclercia, Lelliottia, Shigella)*, Burkholderiaceae *(Burkholderia, Paraburkholderia)*, Pseudomonadaceae *(Pseudomonas)*, Erwiniaceae *(Erwinia. Pantoea)* and Yersiniaceae *(Serratia, Yersinia)* (Figure 4B and Table S1C). Examples of *S. enterica* subsp*. diarizonae* (SPI-20 and SPI-21 T6SSs) and subsp*. houtenae* (SPI-19 T6SS) were detected in Tae7 family.

**Figure 4.**
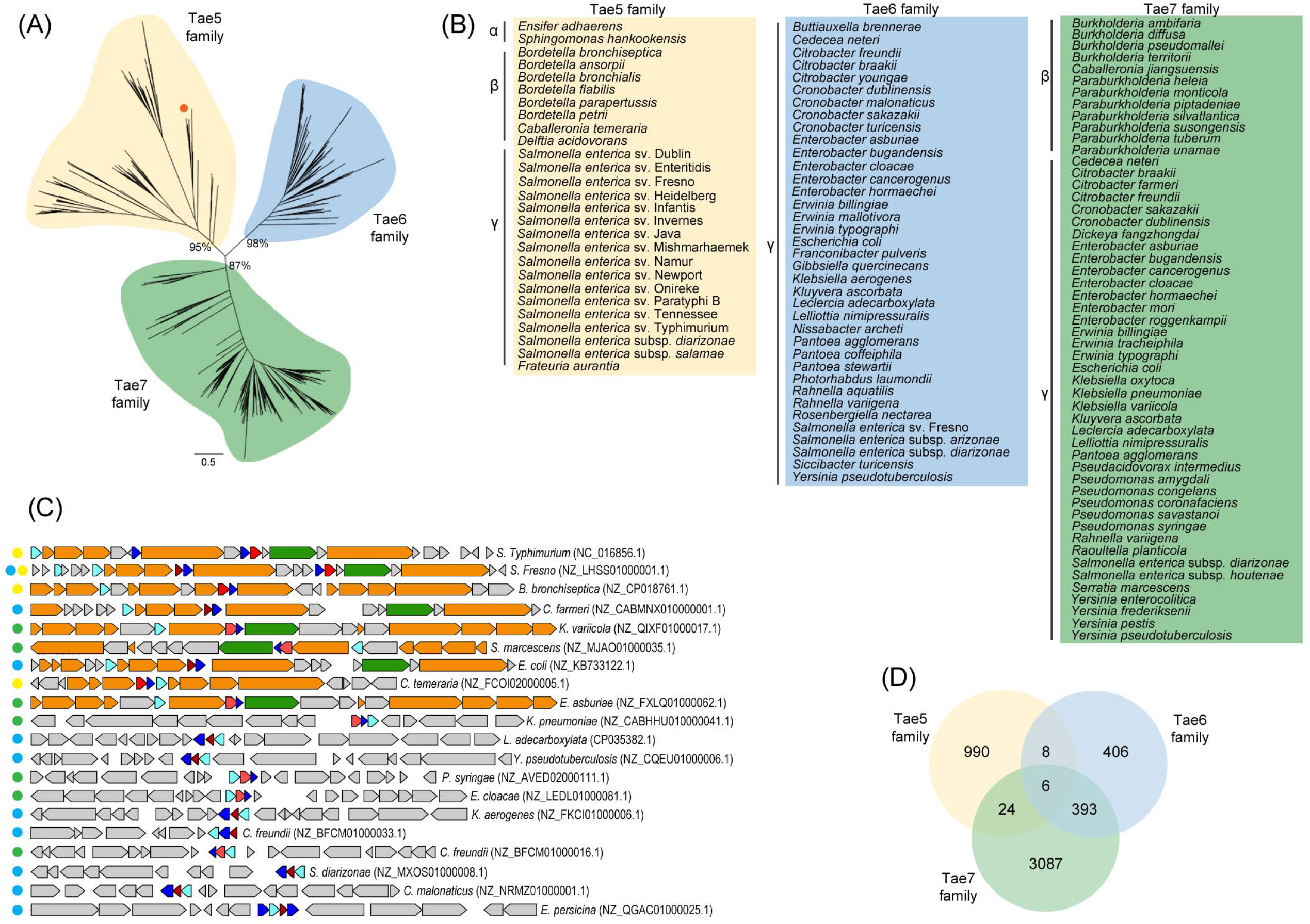
DUF2778 superfamily is widespread and segregates into three families. (A) Maximum likelihood phylogenetic tree of DUF2778 superfamily (clade 1; Figure 3A), showing its division into three families (Tae5, Tae6 and Tae7). Non-parametric bootstrap values at the branches defining each family are indicated. The position of Tae5^STM^ is highlighted by an orange dot. (B) List of common bacterial species encoding members of Tae5, Tae6 and Tae7. The toxic effector is widespread and detected among α-, β- and γ-proteobacteria. (C) Venn diagram representing the total number of bacterial genomes that encode one or more Tae5/6/7 family member. (D) Schematic representation of the genomic context of Tae5/6/7 family members. T6SS structural genes (orange), VgrG genes (green), Hcp genes (light blue), Tae5/6/7 family members (red) and their cognate immunity proteins (dark blue). Bacterial species and genome accession number is described on the right-side of each cluster and colored dots on the left-side denote the family to which the effector belongs: yellow (Tae5), blue (Tae6), green (Tae7).

Examining the genomic context of Tae5-7 family members, we observed that several proteins are encoded within T6SS loci or associated with orphan Hcp proteins (Figure 4C and Table S2), suggesting that these effectors are mainly secreted as cargo associated with Hcp proteins. Figure 4D shows the number of bacterial genomes that have one or more proteins belonging to each of the three families (Tae5-7) (Table S1D). From the total of 1028 genomes containing a Tae5 member, 615 (59.8%) are encoded in the vicinity of a T6SS structural gene (Figure 4C). The same high frequency is observed for Tae6 and Tae7 families with 76.6% (623 out of 813) and 42.6% (1495 out of 3510) of the genomes containing a member in the vicinity of T6SS structural genes, respectively.

Interestingly, genomic analysis revealed that some organisms encode more than one member of the DUF2778 superfamily (Figures 4C and 4D). *S. enterica* serovar Fresno (*S*. Fresno) encodes two members within its SPI-6 T6SS, Tae5^SF^ (WP_000968384.1) and Tae6^SF^ (WP_058115706.1). Tae5^SF^ is closely related to *S*. Typhimurium Tae5^STM^ (STM14_0336), but the Tae6^SF^ homolog is absent in *S*. Typhimurium 14028s (Figure 4C). A close inspection of the *S*. Typhimurium 14028s SPI-6 T6SS locus revealed that it contains an orphan Tai6^STM^ immunity protein (STM14_0332) orthologous to *S*. Fresno Tai6^SF^ (WP_077921582.1) in the same genomic location (Figures 1A and 4C). Curiously this gene is located next to a Tai4 orphan immunity protein homologous to Rap1a (STM14_0331), which confers immunity to the Tae4 family member Ssp1 from *Serratia marcescens* (English et al., 2012). Growth inhibition assays showed that Tai6^STM^ immunity protein (STM14_0332) was not able to neutralize the toxicity of Tae5^STM^ (STM14_0336), thus confirming the specificity of each immunity protein to its toxic effector (Figure S3) and suggesting that Tai6^STM^ and Rap1a may have a role in immunity against effectors secreted by other bacteria.

The domain architecture of most DUF2778-containing proteins is usually quite simple, with most members composed of a single DUF2778 domain; but a few examples such as *Shigella sonnei* WP_052983304.1 present DUF2778 at the C-terminal region of rearrangement hotspot (Rhs) proteins, which are encoded close to PAAR-containing proteins, suggesting secretion via T6SS (Koskiniemi et al., 2013). These Rhs-DUF2778 proteins are mainly represented in Tae7 family in which 11% (387 out of 3510) of the genomes display a member fused to Rhs protein (Figure 4D). Moreover, DUF2778 genes could be found located in the vicinity of genes encoding other secretion system such as T1SS *(abc, omf. mfp*), T2SS *(gspD-N)*, T4SS *(virb1-10)*, T5SS (PF03797, PF03895, translocator) (Table S2) (Abby et al., 2016), suggesting that these DUF2778-containing proteins might have been recruited to be part of polymorphic toxin cassettes that are known to be associated with various protein secretion systems (Zhang et al., 2012). These results are in line with bioinformatic analysis that predicted polymorphic toxins with L,D-peptidase domains fused to Rhs repeats or domains related to secretory systems and were linked to neighboring immunity proteins DUF2750/Imm16 and Imm57 (Zhang et al., 2012).

## Discussion

The peptidoglycan forms a closed, bag-shaped structure surrounding the cytoplasmic membrane and gives bacteria form and resistance to osmotic pressure. It is made of glycan strands crosslinked by short peptides, whose composition is highly diverse among different bacterial species (Vollmer et al., 2008). Such variability in peptidoglycan structure might be the reason why T6SS-containing bacteria use such a diverse array of effectors with amidase activity to target competitors. In this study, we identified and characterized a new superfamily of T6SS amidase effectors, which is evolutionarily related to L,D-transpeptidases containing YkuD domain (formerly called ErfK/YbiS/YcfS/YnhG). The founding member of this superfamily, Tae5^STM^, displays L,D-carboxypeptidase activity and cleaves the bond between *m*DAP^3^ and D-Ala^4^ within the same peptide stem. The specificity of Tae5 is different from the other previously characterized T6SS amidase effectors (Tae1-4) (Russell et al., 2012). Tae1 and Tae4 cleave the bond between D-*i*Glu^2^ and *m*DAP^3^ within the same peptide stem, while Tae2 and Tae3 cleave the crosslink bridge between D-Ala^4^ and *m*DAP^3^ of different peptide stems (Russell et al., 2012). In addition, the evolutionary origin of these two groups of amidase effectors is different: while Tae1-4 catalytic domains are related to CHAP (PF05257) and NlpC/P60 (PF00877) amidases (Anantharaman and Aravind, 2003; Bateman and Rawlings, 2003), Tae5-7 catalytic domains are related to YkuD of L,D-transpeptidases (PF03734) (Biarrotte-Sorin et al., 2006).

L,D-transpeptidases form *m*DAP^3^-*m*DAP^3^ (3 → 3) crosslinks between peptide stems by transferring the peptide bond between the third residue of a tetrapeptide donor stem to the side-chain amide group of a third residue of an adjacent acceptor stem. The catalytic mechanism was proposed to occur in a two-step enzymatic reaction requiring a catalytic cysteine residue; it involves the acylation of the enzyme by the penultimate peptide of the donor stem with the release of the C-terminal amino acid residue (D-Ala^4^), followed by deacylation of this acyl-enzyme intermediate by an acceptor stem (Biarrotte-Sorin et al., 2006; Erdemli et al., 2012). Enzymes displaying L,D-transpeptidase activity (Ldt_Bs_, Ldt_fm_, Ldt_Mt_) (clades 2 and 3; Figure 3A) have an elongated active site with the catalytic triad accessible via two paths. On the other hand, the Csd6 from *H. pylori*, which displays an L,D-transpeptidase domain but only L,D-carboxypeptidase activity (clade 4; Figure 3A), has the active site in a deep pocket with the catalytic triad positioned at the bottom accessible via a single narrow path (Kim et al., 2015). The absence of detailed structural information for Tae5^STM^ does not allow us to determine its precise catalytic mechanism at this point; however, we speculate that structural features that restrict access to its active site might explain its L,D-carboxypeptidase activity. The D-methionine exchange assay showed that Tae5^STM^ is able to cleave the *m*DAP^3^-D-Ala^4^ peptide bond from GM-tetrapeptides and form a new peptide bond between mDAP^3^-D-Met^4^ when D-Met is provided in excess (Figure S2B); however, we are unsure whether this exchange activity is relevant in physiological levels of D-amino acids. We detected a small increase of GM-AE*m*DAP<u>G</u> (13%) in *E. coli* cells expressing SP-Tae5^STM^ compared to cells with empty plasmid (6%) or catalytic inactive SP-Tae5^STM^_C131A_ (6%) (Figure 3E). Such increase could be driven by the exchange activity of SP-Tae5^STM^ or another endogenous enzyme trying to compensate for the reduction of GM-AE*m*DAP<u>A</u>. Together, the enzymatic assays support our hypothesis that the active site of Tae5^STM^ is not accessible to larger substrates such as GM-tetrapeptides and the enzyme preferentially uses small molecules such as water and free amino acids as acceptor substrates.

The toxic phenotype induced by periplasmic expression of Tae5^STM^ seems to be the result of a series of events. First, up to 4.5h of intoxication, cells stop dividing but continue growing in length. It has been shown that cell division is mediated by filaments of FtsZ and FtsA that treadmill circumferentially around the division ring and drive the motion of peptidoglycan-synthesizing enzymes (Bisson-Filho et al., 2017). However, the rate of division septum closure is mainly determined by the D,D-transpeptidase activity of FtsI, indicating that cell wall synthesis plays a limiting role in septum closure (Coltharp et al., 2016). Our hypothesis that Tae5^STM^ promotes toxicity by depleting the pool of acceptor GM-tetrapeptides, thus preventing the formation of new crosslinks by endogenous D,D-transpeptidases, fits with the idea that septum closure rate is limited by peptidoglycan synthesis. Second, the phenotype observed after 4.5h in which cells form blebs and lyse (Movie S2) is also aligned with the idea of diminished peptidoglycan synthesis and crosslink rates. When the peptidoglycan growth rate falls behind that of overall cell growth, the peptidoglycan net stretches resulting in changes in pore size that alters the integrity of the cell wall (Typas et al., 2011).

In T6SS-positive organisms, DUF2778-containing proteins are encoded in bicistrons with their putative immunity protein (Figure 4C). Divergence among immunity proteins was considerably higher than that found within the effector families. The immunity protein for Tae5 family has a DUF2195 domain (Tai5), while Tai6 and Tai7 immunity proteins are encoded by genes with no annotated domain. Tai5 and Tai6 immunity proteins encode a Sec signal peptide for periplasmic localization, while Tai7 immunity proteins have one or more transmembrane domains (Figure S1) (Krogh et al., 2001). A similar diversity in localization signals of T6SS immunity proteins was observed for homologs of the toxin encoded at the C-terminal of VgrG2c (DUF4157) of *Pseudomonas aeruginosa* (Wood et al., 2019). The greater sequence divergence in immunity proteins has been reported previously and is likely due to less restrictive selective pressure compared to effectors; while immunity proteins are selected for effector binding, effectors are selected for both immunity protein binding and catalysis (Russell et al., 2012). Due to the small length of these immunity proteins and their diversity, which impeded automated bioinformatic analysis, we did not attempt further assessment of their phylogeny.

T6SSs translocate effectors by decorating the Hcp-VgrG-PAAR puncturing device. (Cianfanelli et al., 2016; Jana and Salomon, 2019). Small effector proteins composed of only one toxic domain, such as Tae5^STM^, usually interact with Hcp proteins for secretion as their size favors fitting inside the narrow tube formed by Hcp hexamers (Silverman et al., 2013; Jana and Salomon, 2019). Further supporting this notion, our bioinformatic analysis showed members of the three families Tae5, Tae6 and Tae7 associated with orphan Hcp proteins (Figure 4C).

The distribution of Tae5, Tae6 and Tae7 effectors between different organisms give an idea of how these toxins are used by different species to intoxicate competitors. It is interesting to note that most *S. enterica* subsp*. enterica* serovars encode only a Tae5 family member (e.g. *S*. Typhimurium). According to our analysis, Tae6 family members are mainly represented among species of the Enterobacteriaceae *(Cedecea, Citrobacter, Cronobacter, Enterobacter, Escherichia, Franconibacter, Gibbsiella, Klebsiella, Kluyvera, Leclercia, Lelliottia*), indicating that Tae6/Tai6 effector/immunity pairs must be a commonly used ammunition in the gut environment. Therefore, it does not seem to be a great competitive advantage for *S*. Typhimurium to keep a Tae6 effector in its repertoire as many species already living in the gut seem to encode a Tai6 immunity protein. Similarly, despite most *S. enterica* subsp*. enterica* serovars encode only a Tae5 family member, they frequently encode an orphan Tai6 immunity protein, suggesting that although not attacking competitors using Tae6, *Salmonella* are protected from the attack of members of the microbiota encoding a Tae6 effector. Overall, from the perspective of an enteric pathogen that needs to kill competitor species in the gut to colonize the environment, *S. enterica* subsp*. enterica* serovars are well equipped with a Tae5 family member, which is not present in most members of Enterobacteriaceae.

## Methods

### Bacterial strains and growth conditions

All bacterial strains used in this study are listed in Table S3. Strains were grown at 37°C in Lysogeny Broth (10 g/L tryptone, 10 g/L NaCl, 5 g/L yeast extract) under agitation. Cultures were supplemented with antibiotics in the following concentration when necessary: 50 μg/mL kanamycin, 100 μg/mL ampicillin, 50 μg/mL streptomycin, 15 μg/mL chloramphenicol. *S*. Typhimurium mutant strains were constructed by λ-Red recombination engineering using a one-step inactivation procedure (Datsenko and Wanner, 2000).

### Cloning and mutagenesis

All plasmids and primers are listed in Table S3. *STM14_0336* was amplified by PCR and cloned into pBRA vector under the control of P_BAD_ promoter (M. Marroquin, unpublished) with or without PelB signal peptide sequence from pET22b (Novagen) (Bayer-Santos et al., 2019). Immunity proteins *(STM14_0332* and *STM14_0335)* were cloned into pEXT22 under the control of P_TAC_ promoter (Dykxhoorn et al., 1996). For protein expression and purification, STM14_0336 residues between 2-174 were cloned into pET28a (Novagen), including a N-terminal His-tag. Point mutations were created using QuikChange II XL Site-Directed Mutagenesis Kit (Agilent Technologies) and pBRA SP-Tae5^STM^ (STM14_0336) plasmid was used as template. All constructs were confirmed by sequencing.

### Growth inhibition assay

Overnight cultures of *E. coli* DH5α co-expressing effectors for cytoplasmic (pBRA-Tae5^STM^) or periplasmic (pBRA SP-Tae5^STM^) localization and immunity proteins (pEXT22-Tai5^STM^) were serially diluted in LB (1:4) and 5 μL were spotted onto LB-agar (1.5%) containing either 0.2% D-glucose or 0.2% L-arabinose and 200 μM IPTG and incubated at 37°C. Images were acquired after 24h.

### Subcellular fractionation

Subcellular fractionation was adapted from Gauthier et al. (2003) and English et al. (2012) and is based on osmotic shock and ultracentrifugation. Briefly, *E. coli* cells harboring pEXT22-Tai5^STM^-FLAG grown overnight with 200 μM of IPTG were harvested (aliquot corresponding to total proteins was collected for analysis), washed twice with phosphate buffer saline (PBS) and resuspended in 1 mL of 50 mM Tris-HCl pH 7.4, 20% sucrose, 10 mM EDTA and protease inhibitor. Cells were incubated for 10 min at 30°C and recovered by centrifugation (10 min, 8000 *g*, 22°C). Pellets were resuspended in 1 mL of ice-cold water and incubated for 10 min on ice. After centrifugation (10 min, 8000 *g*, 4°C), 900 μL of the supernatant was retained for analysis (enriched in periplasmic proteins). The pellet containing cytoplasmic and membrane proteins was resuspended in 1 mL of sonication buffer (10 mM Tris-HCl pH 7 and protease inhibitor), sonicated 6 rounds of 15 secs with amplitude 30% and centrifuged (15 min, 16000 *g*, 4°C) to remove insoluble proteins. Supernatant was transferred to an ultracentrifuge tube and centrifugated (1h, 50000 *g*, 4°C). Supernatant (900μL) corresponding to the cytoplasmic fraction was retained for analysis, and the pellet with total membrane proteins was washed with sonication buffer once, centrifugated (1h, 50000 *g*, 4°C) and resuspended in SDS-PAGE buffer. Fractions were precipitated with 4 volumes of acetone and resuspended in equivalent volumes of SDS-PAGE buffer. Protein extracts were separated by SDS-PAGE and analyzed by western blot with anti-FLAG (Sigma-Aldrich #F7425), anti-DNAk (Abcam #ab69617), anti-maltose binding protein (MPB) (New England BioLabs #E8032L) and anti-outer membrane protein (OmpA) antibodies.

### Microscopy

For time-lapse microscopy, LB-agar pads were prepared by cutting a rectangular piece out of a double-sided adhesive tape which was taped onto a microscopy slide as described previously (Bayer-Santos et al., 2019). *E. coli* DH5α harboring pBRA SP-Tae5^STM^ was spotted onto 1.5% LB-agar pads supplemented either with 0.2% D-glucose or 0.2% L-arabinose. Phase contrast images were taken every 15 min for 24h using a Zeiss AxioVert.Z1 microscope fitted with an AxioCam MRm camera and an α Plan-Apochromat 63x oil objective. Images were analyzed using FIJI software (Schindelin et al., 2012). To quantify the percentage of dividing or non-dividing cells and the cell doubling time, approximately 100 cells were analyzed in each condition. To determine cell length, approximately 30 cells were measured at each time point (from 30 min to 4.5h), totaling 150 measurements. To visualize membrane labelling, overnight cultures of *E. coli* carrying pBRA SP-Tae5^STM^ were harvested by centrifugation (3 min, 8000 *g*), washed twice with LB and normalized to OD_600nm_ 0.5. Membrane dye FM4-64 (Molecular Probes) were mixed to cells at 1:1 and 4 μL were spotted onto 1.5% LB-agarose pads supplemented with 0.2% D-glucose or 0.2% L-arabinose. After 20h, cells were imaged using a Zeiss AxioVert.A1 microscope fitted with an AxioCam ICm1 camera and a FLUAR 100x oil objective. To analyze FtsZ localization, overnight cultures of *E. coli* FtsZ-mVenus expressing pBRA SP-Tae5^STM^ were diluted 1:100 and grown in liquid LB media to an OD_600nm_ 0.2. Cells were harvested by centrifugation (3 min, 8000 *g*), washed twice with LB and added 0.2% D-glucose or 0.2% L-arabinose. At OD_600nm_ 0.5 cells were harvested, washed twice with PBS and 4 μL were spotted onto 1.5% PBS-agarose pads supplemented with 0.2% D-glucose or 0.2% L-arabinose. Cells were imaged using a Zeiss AxioVert.A1 microscope fitted with an AxioCam ICm1 camera.

### Recombinant protein expression and purification

*E. coli* SHuffle cells expressing pET28a-Tae5^STM^ were subcultured in LB and grown at 37°C to OD_600nm_ 0.7 prior to induction with 0.4 mM IPTG for 16h at 16°C (150 rpm). Cells were harvested by centrifugation, resuspended with buffer (20 mM Tris-HCl pH 7.35, 200 mM NaCl) and lysed by 10 passages in a French Press system. The lysate soluble fraction was loaded onto a 5 mL HiTrap chelating HP column (GE Healthcare) immobilized with 100 mM cobalt chloride and equilibrated with the lysis buffer. After the removal of unbound proteins, the fusion protein was eluted with lysis buffer supplemented with 400 mM imidazole. Purified proteins were concentrated in Amicon Filter Units (Millipore) before purification by size exclusion chromatography using a HiLoad 26/600 Superdex 75 column (GE Healthcare).

### Peptidoglycan purification and enzymatic assays

Peptidoglycan was purified as described by Mesnage et al. (2008) with small modifications. Briefly, bacterial cells were grown in 1 L of LB to OD_600nm_ 0.7 and harvested by centrifugation (15 min, 4000 *g*, 15°C). Pellets were washed with PBS and resuspended in 20 mL of PBS. Cell suspensions were added dropwise to a glass flask containing 80 mL of boiling 5% SDS and incubated for 30 min. Lysates were washed 4 times with water and treated with pronase 2 mg/mL overnight at 60°C. The following day, 1% SDS was added, incubated for 10 min at 95°C and washed 4 times with water. Muropeptides were obtained by digestion with mutanolysin (Sigma #M9901), followed by reduction with sodium borohydride in borate buffer and separation in reverse-phase high performance liquid chromatography (RP-HPLC) (Mesnage et al., 2008). For *in vitro* assays, 10 μM of purified recombinant Tae5^STM^ protein was added to 100 μM of monomeric or dimeric GM-tetrapeptide in buffer containing 50 mM Tris pH 7.5 and 50 mM NaCl and incubated for 4h at 37°C. Digestion products were analyzed by RP-HPLC-MS as described previously (Rodriguez-Rubio et al., 2016). For the amino acid exchange assay, reactions were performed in the same conditions with the addition of 1 mM D-methionine. For the muropeptide profile analysis, *E. coli* carrying empty pBRA, pBRA SP-Tae5^STM^ and pBRA SP-Tae5^STM^_C131A_ were used. Cells were grown overnight, subcultured 1:100 in 250 mL of LB and grown until OD_600nm_ 0.5 with 50 μg/mL streptomycin and 0.2% D-glucose. Cells were washed twice with 20 mL of LB, added 0.2% L-arabinose and incubated for 3h until peptidoglycan was extracted as described above. MS data were analyzed with MassHunter software (Agilent Technologies).

### Bioinformatic analysis

Iterative profile searches using JackHMMER (Potter et al., 2018) with a cutoff evalue of 10^-6^ and a maximum of twenty iteration were performed to search for similar sequences in the non-redundant (nr) protein database from the National Center for Biotechnology Information (NCBI). Similarity-based clustering of proteins was carried out using MMseqs software (Steinegger and Soding, 2017). Sequences alignments were produced with MAFFT local-pair algorithm (Katoh et al., 2005) and non-informative columns were removed with trimAl software (Capella-Gutierrez et al., 2009). Approximately-maximumlikelihood phylogenetic tree were built using FastTree 2 (Price et al., 2010). Proteins were annotated using the Pfam database (El-Gebali et al., 2019) and protein secretion systems were identified using models from TXSSdb (Abby et al., 2016) and the HMMER package (Potter et al., 2018). To collect the neighborhood of the genes of interest an in-house python script was used based on information downloaded from the complete genomes and nucleotide sections of the NCBI database.

## Acknowledgements

We are grateful to Chuck S. Farah (University of Sao Paulo) for sharing equipment, reagents and for careful reading of this manuscript; Marcelo Brocchi (University of Campinas) and Alexandre Bisson-Filho (Brandeis University) for bacterial strains; David Holden (Imperial College London) for antibodies; Mario C. Cruz and Iuri C. Valadão from CEFAP-USP for support with microscopy analysis; Gilberto H. Kaihami for assistance with bioinformatic data collection; and Daniel H. S. Limache for technical support. Mass spectrometry analyses were performed by the chemMS Facility of the Faculty of Science Mass Spectrometry Centre at the University of Sheffield. This work was supported by Sao Paulo Research Foundation (FAPESP) grants to E.B.-S. (2017/02178-2), R.F.S. (2016/09047-8), C.R.G. (2017/17303-7, 2019/00195-2). FAPESP fellowships were awarded to S.S.S. (2018/13819-1), J.T.H. (2018/25316-4), B.Y.M. (2016/00458-5), E.B.-S. (2018/04553-8). G.G.N. was awarded a CAPES fellowship. A.P. is supported by a Biotechnology and Biological Sciences Research Council studentship (BB/M011151/1).

## Author contributions

S.S.S., J.T.H. and E.B.-S. outlined the study. S.S.S., J.T.H., B.Y.M., E.B.-S. performed experiments. S.S.S., J.T.H., and E.B.-S. analyzed data. A.P. and S.M. performed peptidoglycan and enzymatic assays and analyzed the corresponding data. G.G.N. and R.F.S. performed bioinformatic analyses. S.S.S., J.T.H., S.M., G.G.N, R.F.S., C.R.G. and E.B.-S. contributed with scientific discussions. S.S.S. and E.B.-S. wrote the manuscript.

## Declaration of Interests

The authors declare no competing interests.

---

**Movie S1.** Time-lapse microscopy showing *E. coli* cells containing pBRA-Tae5^STM^ grown on LB-agar pads with 0.2% D-glucose (repressed). Images were acquired every 15 min. Scale bar 5 μm. Timestamps in hours:minutes.

**Movie S2.** Time-lapse microscopy showing *E. coli* cells containing pBRA-Tae5^STM^ grown on LB-agar pads with 0.2% L-arabinose (induced). Images were acquired every 15 min. Scale bar 5 μm. Timestamps in hours:minutes.

**Table S1.** Complete list of Tae5, Tae6 and Tae7 family members and list the bacterial genomes that encode one or more Tae5/6/7 family member.

**Table S2.** Complete list of Tae5, Tae6 and Tae7 family members with their genomic context.

**Table S3.** List of bacterial strains, plasmids and primers used in this study.

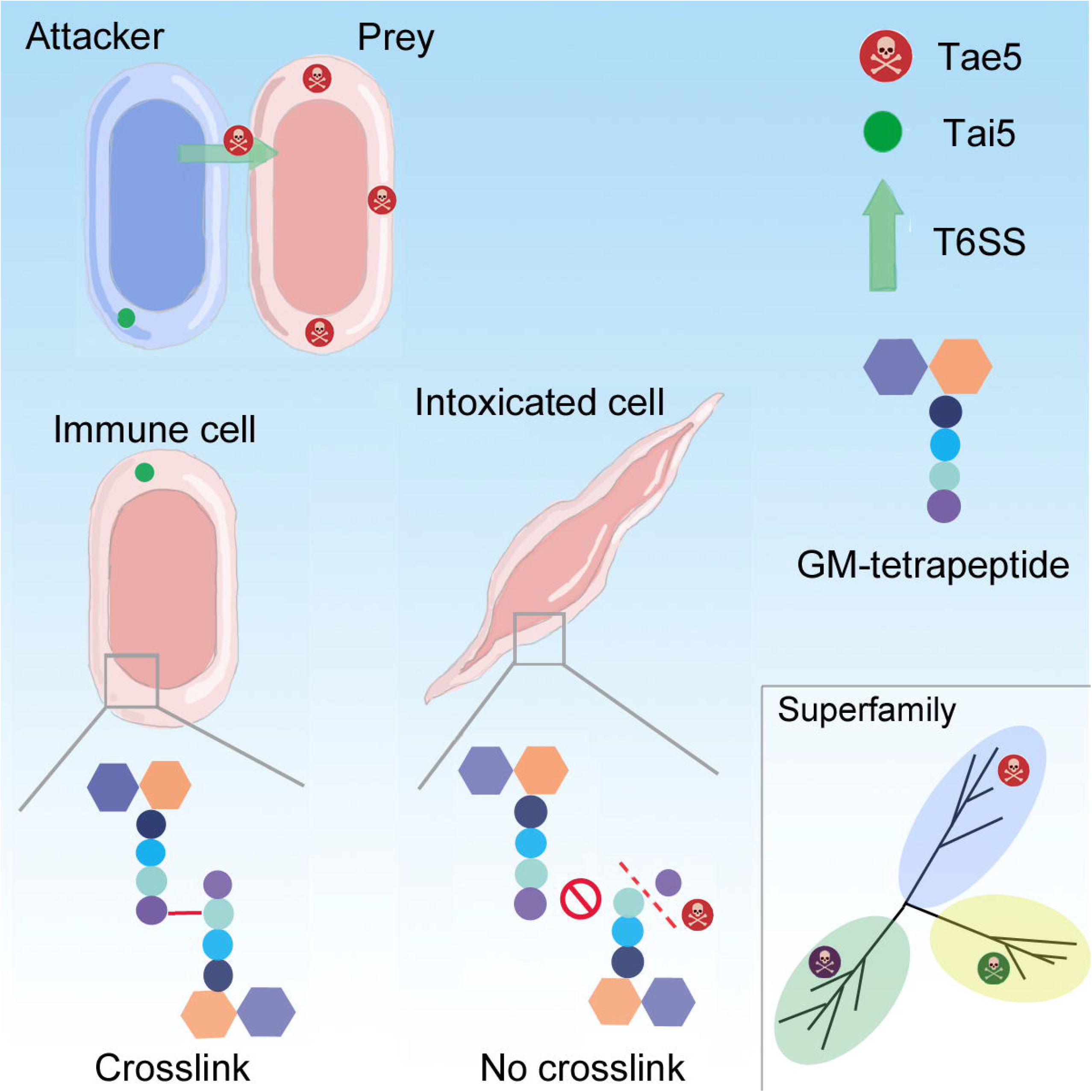

## References

Abby, S.S., Cury, J., Guglielmini, J., Neron, B., Touchon, M. and Rocha, E.P. (2016). Identification of protein secretion systems in bacterial genomes. Sci Rep, 6, 23080.

Ahmad, S., Wang, B., Walker, M.D., Tran, H.R., Stogios, P.J., Savchenko, A., Grant, R.A., Mcarthur, A.G., Laub, M.T. and Whitney, J.C. (2019). An interbacterial toxin inhibits target cell growth by synthesizing (p)ppApp. Nature, 575, 674–678.

Almagro Armenteros, J.J., Tsirigos, K.D., Sonderby, C.K., Petersen, T.N., Winther, O., Brunak, S., Von Heijne, G. and Nielsen, H. (2019). SignalP 5.0 improves signal peptide predictions using deep neural networks. Nat Biotechnol, 37, 420–423.

Anantharaman, V. and Aravind, L. (2003). Evolutionary history, structural features and biochemical diversity of the NlpC/P60 superfamily of enzymes. Genome Biol, 4, R11.

Bao, H., Zhao, J.H., Zhu, S., Wang, S., Zhang, J., Wang, X.Y., Hua, B., Liu, C., Liu, H. and Liu, S.L. (2019). Genetic diversity and evolutionary features of type VI secretion systems in *Salmonella*. Future Microbiol, 14, 139–154.

Bateman, A. and Rawlings, N.D. (2003). The CHAP domain: a large family of amidases including GSP amidase and peptidoglycan hydrolases. Trends Biochem Sci, 28, 2347.

Bayer-Santos, E., Cenens, W., Matsuyama, B.Y., Oka, G.U., Di Sessa, G., Mininel, I.D.V., Alves, T.L. and Farah, C.S. (2019). The opportunistic pathogen *Stenotrophomonas maltophilia* utilizes a type IV secretion system for interbacterial killing. PLoS Pathog, 15, e1007651.

Benz, J., Reinstein, J. and Meinhart, A. (2013). Structural Insights into the Effector - Immunity System Tae4/Tai4 from *Salmonella typhimurium*. PLoS One, 8, e67362.

Bianchet, M.A., Pan, Y.H., Basta, L.a.B., Saavedra, H., Lloyd, E.P., Kumar, P., Mattoo, R., Townsend, C.A. and Lamichhane, G. (2017). Structural insight into the inactivation of *Mycobacterium tuberculosis* non-classical transpeptidase LdtMt2 by biapenem and tebipenem. BMC Biochem, 18, 8.

Biarrotte-Sorin, S., Hugonnet, J.E., Delfosse, V., Mainardi, J.L., Gutmann, L., Arthur, M. and Mayer, C. (2006). Crystal structure of a novel beta-lactam-insensitive peptidoglycan transpeptidase. J Mol Biol, 359, 533–8.

Bielnicki, J., Devedjiev, Y., Derewenda, U., Dauter, Z., Joachimiak, A. and Derewenda, Z.S. (2006). *B. subtilis* ykuD protein at 2.0 A resolution: insights into the structure and function of a novel, ubiquitous family of bacterial enzymes. Proteins, 62, 144–51.

Bisson-Filho, A.W., Hsu, Y.P., Squyres, G.R., Kuru, E., Wu, F., Jukes, C., Sun, Y., Dekker, C., Holden, S., Vannieuwenhze, M.S., et al. (2017). Treadmilling by FtsZ filaments drives peptidoglycan synthesis and bacterial cell division. Science, 355, 739–743.

Blondel, C.J., Jimenez, J.C., Contreras, I. and Santiviago, C.A. (2009). Comparative genomic analysis uncovers 3 novel loci encoding type six secretion systems differentially distributed in *Salmonella* serotypes. BMC Genomics, 10, 354.

Brunet, Y.R., Khodr, A., Logger, L., Aussel, L., Mignot, T., Rimsky, S. and Cascales, E. (2015). H-NS Silencing of the *Salmonella* Pathogenicity Island 6-Encoded Type VI Secretion System Limits *Salmonella enterica* Serovar Typhimurium Interbacterial Killing. Infect Immun, 83, 2738–50.

Capella-Gutierrez, S., Silla-Martinez, J.M. and Gabaldon, T. (2009). trimAl: a tool for automated alignment trimming in large-scale phylogenetic analyses. Bioinformatics, 25, 1972–3.

Caveney, N.A., Caballero, G., Voedts, H., Niciforovic, A., Worrall, L.J., Vuckovic, M., Fonvielle, M., Hugonnet, J.E., Arthur, M. and Strynadka, N.C.J. (2019). Structural insight into YcbB-mediated beta-lactam resistance in *Escherichia coli*. Nat Commun, 10, 1849.

Chen, L., Song, N., Liu, B., Zhang, N., Alikhan, N.F., Zhou, Z., Zhou, Y., Zhou, S., Zheng, D., Chen, M., et al. (2019). Genome-wide Identification and Characterization of a Superfamily of Bacterial Extracellular Contractile Injection Systems. Cell Rep, 29, 511–521 e2.

Cianfanelli, F.R., Monlezun, L. and Coulthurst, S.J. (2016). Aim, Load, Fire: The Type VI Secretion System, a Bacterial Nanoweapon. Trends Microbiol, 24, 51–62.

Coltharp, C., Buss, J., Plumer, T.M. and Xiao, J. (2016). Defining the rate-limiting processes of bacterial cytokinesis. Proc Natl Acad Sci U S A, 113, E1044–53.

Coulthurst, S. (2019). The Type VI secretion system: a versatile bacterial weapon. Microbiology, 165, 503–515.

Datsenko, K.A. and Wanner, B.L. (2000). One-step inactivation of chromosomal genes in *Escherichia coli* K-12 using PCR products. Proc Natl Acad Sci U S A, 97, 6640–5.

Deshazer, D. (2019). A novel contact-independent T6SS that maintains redox homeostasis via Zn(2+) and Mn(2+) acquisition is conserved in the *Burkholderia pseudomallei* complex. Microbiol Res, 226, 48–54.

Dykxhoorn, D.M., St Pierre, R. and Linn, T. (1996). A set of compatible tac promoter expression vectors. Gene, 177, 133–6.

El-Gebali, S., Mistry, J., Bateman, A., Eddy, S.R., Luciani, A., Potter, S.C., Qureshi, M., Richardson, L.J., Salazar, G.A., Smart, A., et al. (2019). The Pfam protein families database in 2019. Nucleic Acids Res, 47, D427–D432.

English, G., Trunk, K., Rao, V.A., Srikannathasan, V., Hunter, W.N. and Coulthurst, S.J. (2012). New secreted toxins and immunity proteins encoded within the Type VI secretion system gene cluster of *Serratia marcescens*. Mol Microbiol, 86, 921–36.

Erdemli, S.B., Gupta, R., Bishai, W.R., Lamichhane, G., Amzel, L.M. and Bianchet, M.A. (2012). Targeting the cell wall of *Mycobacterium tuberculosis:* structure and mechanism of L,D-transpeptidase 2. Structure, 20, 2103–15.

Garcia-Bayona, L. and Comstock, L.E. (2018). Bacterial antagonism in host-associated microbial communities. Science, 361.

Gauthier, A., Puente, J.L. and Finlay, B.B. (2003). Secretin of the enteropathogenic *Escherichia coli* type III secretion system requires components of the type III apparatus for assembly and localization. Infect Immun, 71, 3310–9.

Glauner, B., Holtje, J.V. and Schwarz, U. (1988). The composition of the murein of *Escherichia coli*. J Biol Chem, 263, 10088–95.

Hachani, A., Wood, T.E. and Filloux, A. (2016). Type VI secretion and anti-host effectors. Curr Opin Microbiol, 29, 81–93.

Hood, R.D., Singh, P., Hsu, F., Guvener, T., Carl, M.A., Trinidad, R.R., Silverman, J.M., Ohlson, B.B., Hicks, K.G., Plemel, R.L., et al. (2010). A type VI secretion system of *Pseudomonas aeruginosa* targets a toxin to bacteria. Cell Host Microbe, 7, 25–37.

Jana, B., Fridman, C.M., Bosis, E. and Salomon, D. (2019). A modular effector with a DNase domain and a marker for T6SS substrates. Nat Commun, 10, 3595.

Jana, B. and Salomon, D. (2019). Type VI secretion system: a modular toolkit for bacterial dominance. Future Microbiol, 14, 1451–1463.

Katoh, K., Kuma, K., Toh, H. and Miyata, T. (2005). MAFFT version 5: improvement in accuracy of multiple sequence alignment. Nucleic Acids Res, 33, 511–8.

Kim, H.S., Im, H.N., An, D.R., Yoon, J.Y., Jang, J.Y., Mobashery, S., Hesek, D., Lee, M., Yoo, J., Cui, M., et al. (2015). The Cell Shape-determining Csd6 Protein from *Helicobacter pylori* Constitutes a New Family of L,D-Carboxypeptidase. J Biol Chem, 290, 25103–17.

King, D.T., Sobhanifar, S. and Strynadka, N.C. (2016). One ring to rule them all: Current trends in combating bacterial resistance to the beta-lactams. Protein Sci, 25, 787–803.

Koskiniemi, S., Lamoureux, J.G., Nikolakakis, K.C., T’kint De Roodenbeke, C., Kaplan, M.D., Low, D.A. and Hayes, C.S. (2013). Rhs proteins from diverse bacteria mediate intercellular competition. Proc Natl Acad Sci U S A, 110, 7032–7.

Krogh, A., Larsson, B., Von Heijne, G. and Sonnhammer, E.L. (2001). Predicting transmembrane protein topology with a hidden Markov model: application to complete genomes. J Mol Biol, 305, 567–80.

Kumar, P., Chauhan, V., Silva, J.R.A., Lameira, J., D’andrea, F.B., Li, S.G., Ginell, S.L., Freundlich, J.S., Alves, C.N., Bailey, S., et al. (2017). *Mycobacterium abscessus* l,d-Transpeptidases Are Susceptible to Inactivation by Carbapenems and Cephalosporins but Not Penicillins. Antimicrob Agents Chemother, 61.

Ma, J., Sun, M., Pan, Z., Lu, C. and Yao, H. (2018). Diverse toxic effectors are harbored by vgrG islands for interbacterial antagonism in type VI secretion system. Biochim Biophys Acta Gen Subj, 1862, 1635–1643.

Ma, L.S., Hachani, A., Lin, J.S., Filloux, A. and Lai, E.M. (2014). *Agrobacterium tumefaciens* deploys a superfamily of type VI secretion DNase effectors as weapons for interbacterial competition in planta. Cell Host Microbe, 16, 94–104.

Mainardi, J.L., Fourgeaud, M., Hugonnet, J.E., Dubost, L., Brouard, J.P., Ouazzani, J., Rice, L.B., Gutmann, L. and Arthur, M. (2005). A novel peptidoglycan cross-linking enzyme for a beta-lactam-resistant transpeptidation pathway. J Biol Chem, 280, 38146–52.

Mariano, G., Trunk, K., Williams, D.J., Monlezun, L., Strahl, H., Pitt, S.J. and Coulthurst, S.J. (2019). A family of Type VI secretion system effector proteins that form ion-selective pores. Nat Commun, 10, 5484.

Mesnage, S., Chau, F., Dubost, L. and Arthur, M. (2008). Role of N-acetylglucosaminidase and N-acetylmuramidase activities in *Enterococcus faecalis* peptidoglycan metabolism. J Biol Chem, 283, 19845–53.

Moore, D.A., Whatley, Z.N., Joshi, C.P., Osawa, M. and Erickson, H.P. (2017). Probing for Binding Regions of the FtsZ Protein Surface through Site-Directed Insertions: Discovery of Fully Functional FtsZ-Fluorescent Proteins. J Bacteriol, 199.

Mougous, J.D., Cuff, M.E., Raunser, S., Shen, A., Zhou, M., Gifford, C.A., Goodman, A.L., Joachimiak, G., Ordonez, C.L., Lory, S., et al. (2006). A virulence locus of *Pseudomonas aeruginosa* encodes a protein secretion apparatus. Science, 312, 152–630.

Mulder, D.T., Cooper, C.A. and Coombes, B.K. (2012). Type VI secretion system-associated gene clusters contribute to pathogenesis of *Salmonella enterica* serovar Typhimurium. Infect Immun, 80, 1996–2007.

Nguyen, V.S., Douzi, B., Durand, E., Roussel, A., Cascales, E. and Cambillau, C. (2018). Towards a complete structural deciphering of Type VI secretion system. Curr Opin Struct Biol, 49, 77–84.

Parsons, D.A. and Heffron, F. (2005). sciS, an icmF homolog in *Salmonella enterica* serovar Typhimurium, limits intracellular replication and decreases virulence. Infect Immun, 73, 4338–45.

Potter, S.C., Luciani, A., Eddy, S.R., Park, Y., Lopez, R. and Finn, R.D. (2018). HMMER web server: 2018 update. Nucleic Acids Res, 46, W200–W204.

Price, M.N., Dehal, P.S. and Arkin, A.P. (2010). FastTree 2--approximately maximum likelihood trees for large alignments. PLoS One, 5, e9490.

Renault, M.G., Zamarreno Beas, J., Douzi, B., Chabalier, M., Zoued, A., Brunet, Y.R., Cambillau, C., Journet, L. and Cascales, E. (2018). The gp27-like Hub of VgrG Serves as Adaptor to Promote Hcp Tube Assembly. J Mol Biol, 430, 3143–3156.

Rodriguez-Rubio, L., Gerstmans, H., Thorpe, S., Mesnage, S., Lavigne, R. and Briers, Y. (2016). DUF3380 Domain from a *Salmonella* Phage Endolysin Shows Potent N-Acetylmuramidase Activity. Appl Environ Microbiol, 82, 4975–81.

Russell, A.B., Singh, P., Brittnacher, M., Bui, N.K., Hood, R.D., Carl, M.A., Agnello, D.M., Schwarz, S., Goodlett, D.R., Vollmer, W., et al. (2012). A widespread bacterial type VI secretion effector superfamily identified using a heuristic approach. Cell Host Microbe, 11, 538–49.

Salih, O., He, S., Planamente, S., Stach, L., Macdonald, J.T., Manoli, E., Scheres, S.H.W., Filloux, A. and Freemont, P.S. (2018). Atomic Structure of Type VI Contractile Sheath from *Pseudomonas aeruginosa*. Structure, 26, 329–336 e3.

Sana, T.G., Flaugnatti, N., Lugo, K.A., Lam, L.H., Jacobson, A., Baylot, V., Durand, E., Journet, L., Cascales, E. and Monack, D.M. (2016). *Salmonella* Typhimurium utilizes a T6SS-mediated antibacterial weapon to establish in the host gut. Proc Natl Acad Sci U S A, 113, E5044–51.

Schindelin, J., Arganda-Carreras, I., Frise, E., Kaynig, V., Longair, M., Pietzsch, T., Preibisch, S., Rueden, C., Saalfeld, S., Schmid, B., et al. (2012). Fiji: an open-source platform for biological-image analysis. Nat Methods, 9, 676–82.

Shneider, M.M., Buth, S.A., Ho, B.T., Basler, M., Mekalanos, J.J. and Leiman, P.G. (2013). PAAR-repeat proteins sharpen and diversify the type VI secretion system spike. Nature, 500, 350–353.

Si, M., Wang, Y., Zhang, B., Zhao, C., Kang, Y., Bai, H., Wei, D., Zhu, L., Zhang, L., Dong, T.G., et al. (2017a). The Type VI Secretion System Engages a Redox-Regulated Dual-Functional Heme Transporter for Zinc Acquisition. Cell Rep, 20, 949–959.

Si, M., Zhao, C., Burkinshaw, B., Zhang, B., Wei, D., Wang, Y., Dong, T.G. and Shen, X. (2017b). Manganese scavenging and oxidative stress response mediated by type VI secretion system in *Burkholderia thailandensis*. Proc Natl Acad Sci U S A, 114, E2233–E2242.

Silverman, J.M., Agnello, D.M., Zheng, H., Andrews, B.T., Li, M., Catalano, C.E., Gonen, T. and Mougous, J.D. (2013). Haemolysin coregulated protein is an exported receptor and chaperone of type VI secretion substrates. Mol Cell, 51, 584–93.

Steinegger, M. and Soding, J. (2017). MMseqs2 enables sensitive protein sequence searching for the analysis of massive data sets. Nat Biotechnol, 35, 1026–1028.

Tang, J.Y., Bullen, N.P., Ahmad, S. and Whitney, J.C. (2018). Diverse NADase effector families mediate interbacterial antagonism via the type VI secretion system. J Biol Chem, 293, 1504–1514.

Ting, S.Y., Bosch, D.E., Mangiameli, S.M., Radey, M.C., Huang, S., Park, Y.J., Kelly, K.A., Filip, S.K., Goo, Y.A., Eng, J.K., et al. (2018). Bifunctional Immunity Proteins Protect Bacteria against FtsZ-Targeting ADP-Ribosylating Toxins. Cell, 175, 1380–1392 e14.

Trunk, K., Coulthurst, S.J. and Quinn, J. (2019). A New Front in Microbial Warfare-Delivery of Antifungal Effectors by the Type VI Secretion System. J Fungi (Basel), 5.

Typas, A., Banzhaf, M., Gross, C.A. and Vollmer, W. (2011). From the regulation of peptidoglycan synthesis to bacterial growth and morphology. Nat Rev Microbiol, 10, 123–36.

Vollmer, W. and Bertsche, U. (2008). Murein (peptidoglycan) structure, architecture and biosynthesis in *Escherichia coli*. Biochim Biophys Acta, 1778, 1714–34.

Vollmer, W., Blanot, D. and De Pedro, M.A. (2008). Peptidoglycan structure and architecture. FEMS Microbiol Rev, 32, 149–67.

Wang, J., Brackmann, M., Castano-Diez, D., Kudryashev, M., Goldie, K.N., Maier, T., Stahlberg, H. and Basler, M. (2017). Cryo-EM structure of the extended type VI secretion system sheath-tube complex. Nat Microbiol, 2, 1507–1512.

Wang, J., Yang, B., Leier, A., Marquez-Lago, T.T., Hayashida, M., Rocker, A., Zhang, Y., Akutsu, T., Chou, K.C., Strugnell, R.A., et al. (2018). Bastion6: a bioinformatics approach for accurate prediction of type VI secreted effectors. Bioinformatics, 34, 2546–2555.

Wang, S., Yang, D., Wu, X., Yi, Z., Wang, Y., Xin, S., Wang, D., Tian, M., Li, T., Qi, J., et al. (2019). The Ferric Uptake Regulator Represses Type VI Secretion System Function by Binding Directly to the clpV Promoter in Salmonella enterica Serovar Typhimurium. Infect Immun, 87.

Wang, T., Si, M., Song, Y., Zhu, W., Gao, F., Wang, Y., Zhang, L., Zhang, W., Wei, G., Luo, Z.Q., et al. (2015). Type VI Secretion System Transports Zn2+ to Combat Multiple Stresses and Host Immunity. PLoS Pathog, 11, e1005020.

Whitney, J.C., Chou, S., Russell, A.B., Biboy, J., Gardiner, T.E., Ferrin, M.A., Brittnacher, M., Vollmer, W. and Mougous, J.D. (2013). Identification, structure, and function of a novel type VI secretion peptidoglycan glycoside hydrolase effector-immunity pair. J Biol Chem, 288, 26616–24.

Wood, T.E., Howard, S.A., Forster, A., Nolan, L.M., Manoli, E., Bullen, N.P., Yau, H.C.L., Hachani, A., Hayward, R.D., Whitney, J.C., et al. (2019). The *Pseudomonas aeruginosa* T6SS Delivers a Periplasmic Toxin that Disrupts Bacterial Cell Morphology. Cell Rep, 29, 187–201 e7.

Zhang, D., De Souza, R.F., Anantharaman, V., Iyer, L.M. and Aravind, L. (2012). Polymorphic toxin systems: Comprehensive characterization of trafficking modes, processing, mechanisms of action, immunity and ecology using comparative genomics. Biol Direct, 7, 18.

Zhang, H., Zhang, H., Gao, Z.Q., Wang, W.J., Liu, G.F., Xu, J.H., Su, X.D. and Dong, Y.H. (2013). Structure of the type VI effector-immunity complex (Tae4-Tai4) provides novel insights into the inhibition mechanism of the effector by its immunity protein. J Biol Chem, 288, 5928–39.

